# MCD Stitcher: An open-source tool for whole-slide stitching and conversion of Imaging Mass Cytometry data

**DOI:** 10.64898/2026.06.26.732348

**Authors:** Pawan Chaurasia

## Abstract

Imaging Mass Cytometry (IMC) combines metal-tagged antibody labelling with laser ablation mass spectrometry to generate highly multiplexed spatial images of tissue sections. However, the area that can be acquired within a single region of interest (ROI) is limited by hardware and software constraints, requiring large tissues to be imaged as multiple tiled ROIs. Reconstructing these ROIs into whole-slide images requires additional processing, while the proprietary .mcd file format can hinder integration with standard bioimage analysis workflows.

Here, we present MCD Stitcher, an open-source Python package for converting .mcd files into OME-TIFF images with automated whole-slide stitching. The tool supports rectangular and polygonal ROIs, accommodates variable pixel sizes between ROIs, and uses memory-aware chunked reading during data ingestion to process large datasets on standard workstations. The generated OME-TIFF outputs preserve spatial, channel, and acquisition metadata for downstream analysis in tools such as QuPath, napari, and ImageJ/Fiji.

MCD Stitcher provides a reproducible workflow for converting raw IMC data into interoperable image formats, enabling whole-slide spatial analysis without reliance on vendor-specific software.

## Introduction

Imaging Mass Cytometry (IMC) has emerged as a powerful approach for spatial proteomics, enabling simultaneous detection of 40 or more protein markers at cellular and subcellular resolution across tissue sections.(1,2) As spatial profiling technologies become increasingly important for computational pathology and multimodal tissue analysis, IMC provides the advantage of preserving tissue morphology while capturing high-dimensional molecular information. Commercial platforms from Standard BioTools, including Hyperion, Hyperion Plus, and XTi, generate multiplexed images that reveal tissue architecture, cell–cell interactions, and the spatial organisation of immune populations.(3,4) These capabilities are critical for studying tumour microenvironments, identifying therapeutic targets, and characterising treatment responses.(5,6)

A single acquisition on Standard BioTools IMC systems may contain one or more regions of interest (ROIs), each corresponding to a spatially defined area selected for imaging. In practice, the usable size of an individual ROI is limited by hardware and software constraints, including stage travel range, laser ablation and scanning speed, memory usage, acquisition duration, and software stability. Large tissue sections therefore often need to be acquired as multiple adjacent or partially overlapping ROIs, particularly when tissue-scale coverage is required.

As a result, tissue-scale IMC analysis depends on tiled acquisition followed by downstream stitching into a whole-slide image. This reconstruction step is not trivial: it must preserve spatial coordinates, ROI geometry, variable pixel sizes, and acquisition metadata while also producing output that can be used in standard bioimage analysis workflows. Previous work has shown the value of reconstructing whole-slide IMC images by registering IMC fields of view to a whole-slide immunofluorescence reference image.(7) More broadly, the growing emphasis on FAIR bioimaging data and interoperable formats has highlighted the importance of open standards such as OME-TIFF and emerging OME-NGFF-based approaches.(8,9) However, the proprietary .mcd file format still presents a barrier to direct integration with open bioimage analysis ecosystems, including QuPath, napari, and ImageJ/Fiji.(10–12)

Several open-source tools already address parts of this problem. The imctools package and the steinbock framework convert individual IMC acquisitions to OME-TIFF or TIFF, and the readimc library and the napari-imc plugin enable direct reading of .mcd files.(13) These tools, however, operate at the level of individual acquisitions and do not reconstruct whole-slide images from multiple ROIs, particularly when ROIs are polygonal or have been acquired at different pixel sizes. To address this specific gap, we developed MCD Stitcher, an open-source Python package that converts .mcd files into OME-TIFF images with automated whole-slide stitching. Built upon the readimc library,(13) which provides low-level access to IMC data structures, MCD Stitcher is designed to: (i) support rectangular and polygonal ROIs with variable pixel sizes, (ii) operate efficiently on standard hardware through memory-aware chunked reading and streaming writes, and (iii) preserve OME-XML metadata for compatibility with the broader bioimage analysis ecosystem.

## Results

MCD Stitcher provides four command-line tools that together support the workflow from raw .mcd files to analysis-ready OME-TIFF outputs. A single all-in-one command, `mcd_process`, opens each .mcd file once and performs any combination of operations (per-ROI conversion, whole-slide stitching, panorama export, ROI-to-panorama coordinate mapping, metadata inspection, channel filtering, and pyramid generation) in a single pass, and can process either a single .mcd file or a directory of files for batch workflows. Three focused tools (`mcd_stitch, mcd_convert,`and `tiff_subset`) expose the core stitching, conversion, and post-processing operations individually. All tools write OME-TIFF as either 16-bit unsigned integer or 32-bit float and support optional LZW or zstd compression.

**Whole-slide stitching** is performed using `mcd_stitch`(or `mcd_process –stitch`), which reconstructs a stitched OME-TIFF from ROIs contained within one or more acquisitions. The tool extracts ROI coordinates, timestamps, pixel sizes, and channel labels, then places each ROI onto a global canvas (spanning the bounding box of the selected ROIs) using the recorded spatial coordinates. Polygonal, non-rectangular ROIs are supported through ROI-specific masks, which prevent pixels outside the acquired region from being incorrectly included during compositing. When ROIs have different pixel sizes, each ROI is rescaled to a common global pixel size using bilinear interpolation before being placed onto the stitched canvas. Users can also selectively stitch a subset of ROIs rather than processing all ROIs in the input file.

**Per-ROI conversion** is performed using `mcd_convert`(or `mcd_process--convert`), which exports individual ROIs from an .mcd file as separate OME-TIFF images while preserving associated metadata, including acquisition timestamps and descriptions. This mode is useful for tissue microarray samples or other workflows in which ROIs are analysed individually rather than reconstructed into a larger whole-slide image.

### Panorama export and ROI mapping

During acquisition, the instrument records one or more panorama overview images of the slide together with the spatial coordinates of each ROI. MCD Stitcher can export these panoramas as PNG images, with ROI outlines drawn onto larger panoramas, and can write a text file mapping each ROI to its pixel coordinates within a chosen panorama (`mcd_process --panorama and --roi_map`). This provides a spatial reference that links the acquired ROIs back to the overall slide, which is useful both for documenting the acquisition layout and for relating stitched outputs to the original tissue.

**Channel filtering and pyramid generation** are performed using `tiff_subset`(also available within `mcd_process`via `--filter and --pyramid`), which enables users to reduce file size, remove unnecessary channels, and generate pyramidal OME-TIFF outputs for efficient viewing. Raw stitched or per-ROI OME-TIFF files may include control channels, calibration markers, or autofluorescence channels that are not required for downstream analysis. `tiff_subset`allows selected channels to be retained, compression to be applied, and multi-resolution pyramids to be generated for viewing in platforms such as QuPath and napari, while preserving OME-XML metadata.

The implementation uses a memory-aware processing pipeline. For each acquisition, the raw pixel data are read directly from the readimc-opened .mcd file in chunks of up to 50,000 pixels, in which each pixel is a record containing its X and Y coordinates together with the intensity values for all channels, and each chunk is scattered into the acquisition image array before the next chunk is read, which bounds the peak memory required to parse the raw data stream. During stitching, each ROI is read, rescaled if required, masked to preserve polygonal boundaries, composited onto the global canvas, and then released from memory. This design reduces memory usage during data ingestion and enables large IMC datasets to be processed on standard workstations. The reader also validates that each acquisition’s stored data size is an exact multiple of its per-pixel record size; if this check fails, indicating truncated or corrupted data, it automatically retries in a recovery mode that reads the largest whole number of complete pixels and discards the incomplete remainder. All processing modes can operate on a single .mcd file, and `mcd_process`additionally supports directories of .mcd files for batch processing.

**Fig. 1.**
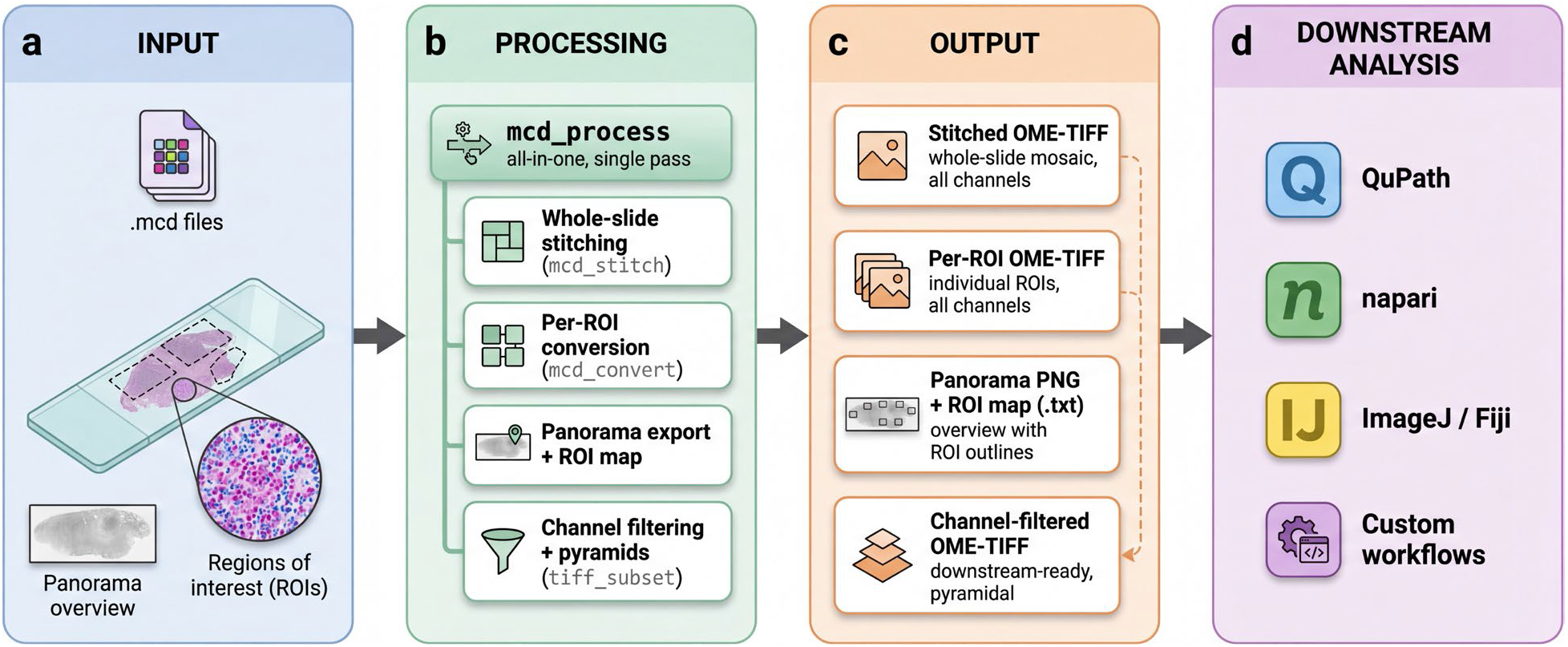
Overview of the MCD Stitcher workflow. (a) IMC data are provided as .mcd files containing multiple tissue regions of interest. (b) MCD Stitcher provides command-line utilities to convert .mcd acquisitions into per-ROI OME-TIFF files, stitch selected ROIs into whole-slide mosaics, and generate channel-filtered pyramid images. (c) The workflow produces standardised OME-TIFF outputs, including stitched mosaics, individual ROI images, and channel-filtered multiscale images. (d) These outputs are compatible with downstream bioimage analysis platforms such as QuPath, napari, ImageJ/Fiji, and custom analysis workflows.

### Demonstration

To demonstrate MCD Stitcher on a representative dataset, we applied it to a tiled IMC acquisition comprising 8 partially overlapping regions of interest acquired with a 36-marker panel at 1 µm pixel size on a Standard BioTools XTi system. Using `mcd_stitch,`the individual ROIs were reconstructed into a single whole-slide OME-TIFF based on their recorded stage coordinates, spanning a 5,105 × 7,074 pixel (36.1-megapixel) canvas. The reconstruction preserved continuous tissue architecture across adjacent tiles (Fig. 2) and retained OME-XML spatial and channel metadata, opening directly in QuPath and napari for downstream analysis.

**Fig. 2.**
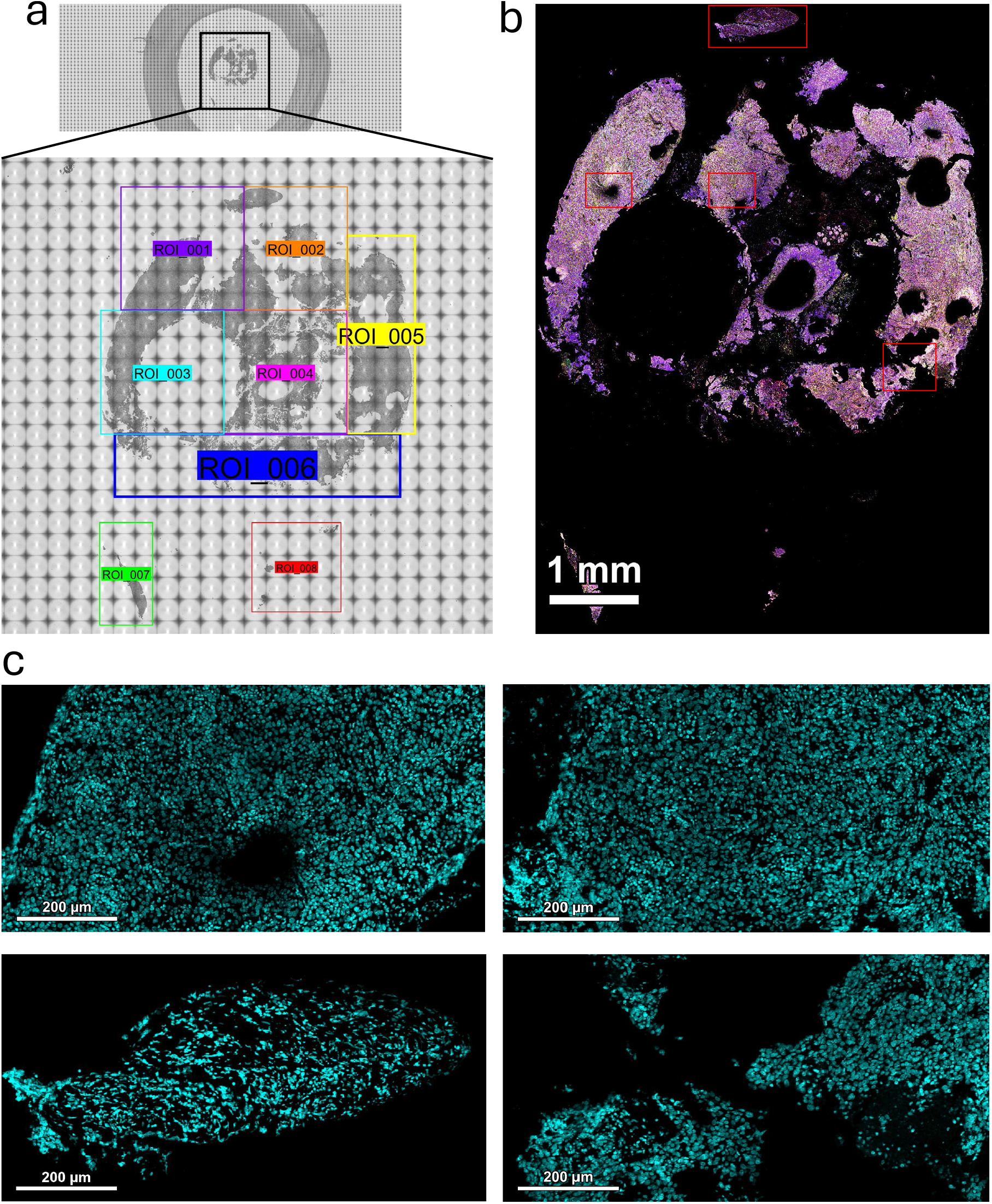
Whole-slide reconstruction of a tiled IMC acquisition. (a) Slide panorama with the outlines of the 8 acquired ROIs overlaid, showing the tiling layout across the tissue core. (b) Whole-slide OME-TIFF reconstructed by MCD Stitcher from the 8 ROIs using their recorded stage coordinates, showing all 36 marker channels with default QuPath channel thresholding. (c) DNA-only magnified example regions showing preserved single-cell resolution across the boundaries between adjacent tiles. Dataset: tiled IMC acquisition, 36-marker panel, 1 µm pixel size, Standard BioTools XTi.

## Discussion

MCD Stitcher addresses a practical gap in IMC workflows. While existing open-source tools can convert individual .mcd acquisitions to OME-TIFF, none provide automated whole-slide reconstruction of multi-ROI acquisitions, including polygonal ROIs and ROIs acquired at different pixel sizes. By integrating whole-slide stitching, per-ROI conversion, panorama export, channel filtering, and pyramid generation within a single installable package, the tool enables researchers to move from raw instrument output to analysis-ready data without reliance on vendor software or custom conversion pipelines.

The choice of OME-TIFF as the output format is deliberate. OME-TIFF is widely supported by bioimage analysis platforms, including QuPath, napari, ImageJ/Fiji, and CellProfiler, and allows spatial and channel metadata to be preserved through embedded OME-XML. This is particularly important for IMC datasets, where physical pixel dimensions, channel labels, and ROI coordinates are required for quantitative spatial analysis. Simpler image formats such as plain TIFF or PNG do not reliably preserve this information.

A key technical challenge in IMC data processing is that acquisitions may contain multiple ROIs with irregular shapes and variable pixel sizes. Many conventional image stitching workflows assume rectangular tiles arranged on a regular grid. In contrast, MCD Stitcher models ROIs as independent spatial entities, enabling reconstruction of whole-slide images from polygonal ROIs and heterogeneous pixel sizes. By combining ROI-specific masking with rescaling to a common coordinate system, the tool provides a stitching workflow tailored to the structure of IMC-derived datasets. MCD Stitcher prioritises reconstruction based on instrument-defined spatial coordinates rather than image-derived registration, reducing dependence on feature matching and preserving acquisition-defined geometry.

By converting raw IMC data into open and widely supported image formats, MCD Stitcher facilitates reproducible and scalable spatial analysis workflows across research and clinical applications.

## Limitations

MCD Stitcher is designed specifically for Standard BioTools Hyperion, Hyperion Plus, and XTi data stored in the .mcd format. It does not currently support other multiplexed imaging platforms, such as CODEX, Xenium, or CosMx, nor alternative file formats from these systems. The stitching workflow assumes that ROIs belong to a common tissue coordinate space and does not perform cross-slide registration or alignment of separate tissue sections.

Memory usage scales with channel count, pixel data type, and stitched tissue area. Although chunked reading reduces peak memory usage during data ingestion, the full output canvas must currently be held in memory during compositing. Consequently, stitched reconstructions spanning tissue areas larger than approximately 1 cm × 1 cm, the nominal upper limit of a single IMC ROI, may require substantial system memory, particularly when many channels are included. In practice, users working with large tissue sections may therefore prefer to process data using the per-ROI conversion workflow or divide the reconstruction into smaller spatial regions.

The current implementation does not include automated overlap detection, seam blending, or intensity normalisation across ROIs. When adjacent ROIs exhibit intensity variation, for example due to laser power drift, staining variation, or tissue thickness differences, these differences may remain visible in the stitched output. Future work may incorporate post-stitching normalisation and blending approaches to address these effects.

## Code availability

MCD Stitcher is available as open-source software under the GNU General Public License v3.0. The source code, documentation, and example usage are available on GitHub (https://github.com/PawanChaurasia/mcd_stitcher). It is implemented in Python (supporting Python 3.9–3.13) and provides both a command-line interface and a Python API. This paper describes MCD Stitcher version 2.3.0.

## Data availability

The IMC dataset used to demonstrate whole-slide stitching is available from the corresponding author on reasonable request, subject to institutional data-sharing agreements.

## Acknowledgements

This research received no specific grant from any funding agency in the public, commercial, or not-for-profit sectors.

## References

1. Giesen C, Wang HAO, Schapiro D, Zivanovic N, Jacobs A, Hattendorf B, et al. Highly multiplexed imaging of tumor tissues with subcellular resolution by mass cytometry. Nat Methods. 2014 Apr;11(4):417–22. doi:10.1038/nmeth.2869

2. Chang Q, Ornatsky OI, Siddiqui I, Loboda A, Baranov VI, Hedley DW. Imaging Mass Cytometry. Cytometry A. 2017;91(2):160–9. doi:10.1002/cyto.a.23053

3. Ali HR, Jackson HW, Zanotelli VRT, Danenberg E, Fischer JR, Bardwell H, et al. Imaging mass cytometry and multiplatform genomics define the phenogenomic landscape of breast cancer. Nat Cancer. 2020 Feb;1(2):163–75. doi:10.1038/s43018-020-0026-6

4. Wang XQ, Danenberg E, Huang CS, Egle D, Callari M, Bermejo B, et al. Spatial predictors of immunotherapy response in triple-negative breast cancer. Nature. 2023 Sep;621(7980):868–76. doi:10.1038/s41586-023-06498-3

5. Kuett L, Catena R, Özcan A, Plüss A, Schraml P, Moch H, et al. Three-dimensional imaging mass cytometry for highly multiplexed molecular and cellular mapping of tissues and the tumor microenvironment. Nat Cancer. 2022 Jan;3(1):122–33. doi:10.1038/s43018-021-00301-w

6. Jackson HW, Fischer JR, Zanotelli VRT, Ali HR, Mechera R, Soysal SD, et al. The single-cell pathology landscape of breast cancer. Nature. 2020 Feb;578(7796):615–20. doi:10.1038/s41586-019-1876-x

7. Kim EN, Chen PZ, Bressan D, Tripathi M, Miremadi A, Pietro M di, et al. Dual-modality imaging of immunofluorescence and imaging mass cytometry for whole-slide imaging and accurate segmentation. Cell Rep Methods. 2023 Oct 23;3(10). doi:10.1016/j.crmeth.2023.100595

8. Goldberg IG, Allan C, Burel JM, Creager D, Falconi A, Hochheiser H, et al. The Open Microscopy Environment (OME) Data Model and XML file: open tools for informatics and quantitative analysis in biological imaging. Genome Biol. 2005;6(5):R47. doi:10.1186/gb-2005-6-5-r47

9. Moore J, Allan C, Besson S, Burel JM, Diel E, Gault D, et al. OME-NGFF: a next-generation file format for expanding bioimaging data-access strategies. Nat Methods. 2021 Dec;18(12):1496–8. doi:10.1038/s41592-021-01326-w

10. Sofroniew N, Lambert T, Bokota G, Nunez-Iglesias J, Sobolewski P, Sweet A, et al. napari: a multi-dimensional image viewer for Python [Internet]. Zenodo; 2026 [cited 2026 May 7]. Available from: https://zenodo.org/records/20058236 doi:10.5281/zenodo.20058236

11. Bankhead P, Loughrey MB, Fernández JA, Dombrowski Y, McArt DG, Dunne PD, et al. QuPath: Open source software for digital pathology image analysis. Sci Rep. 2017 Dec 4;7(1):16878. doi:10.1038/s41598-017-17204-5

12. Schindelin J, Arganda-Carreras I, Frise E, Kaynig V, Longair M, Pietzsch T, et al. Fiji: an open-source platform for biological-image analysis. Nat Methods. 2012 Jun 28;9(7):676–82. doi:10.1038/nmeth.2019

13. Windhager J, Zanotelli VRT, Schulz D, Meyer L, Daniel M, Bodenmiller B, et al. An end-to-end workflow for multiplexed image processing and analysis. Nat Protoc. 2023 Nov;18(11):3565–613. doi:10.1038/s41596-023-00881-0

